# The neutralization effect of Montelukast on SARS-CoV-2 is shown by multiscale *in silico* simulations and combined *in vitro* studies

**DOI:** 10.1101/2020.12.26.424423

**Authors:** Serdar Durdagi, Timucin Avsar, Muge Didem Orhan, Muge Serhatli, Bertan Koray Balcioglu, Hasan Umit Ozturk, Alisan Kayabolen, Yuksel Cetin, Seyma Aydinlik, Tugba Bagci-Onder, Saban Tekin, Hasan Demirci, Mustafa Guzel, Atilla Akdemir, Seyma Calis, Lalehan Oktay, Ilayda Tolu, Yasar Enes Butun, Ece Erdemoglu, Alpsu Olkan, Nurettin Tokay, Şeyma Işık, Aysenur Ozcan, Elif Acar, Sehriban Buyukkilic, Yesim Yumak

## Abstract

Small molecule inhibitors have previously been investigated in different studies as possible therapeutics in the treatment of SARS-CoV-2. In the current drug repurposing study, we identified the leukotriene (D4) receptor antagonist Montelukast as a novel agent that simultaneously targets two important drug targets of SARS-CoV-2. We initially demonstrated the dual inhibition profile of Montelukast through multiscale molecular modeling studies. Next, we characterized its effect on both targets by different *in vitro* experiments including the Fluorescent Resonance Energy Transfer (FRET)-based main protease enzyme inhibition assay, surface plasmon resonance (SPR) spectroscopy, pseudovirus neutralization on HEK293T / hACE2, and virus neutralization assay using xCELLigence MP real time cell analyzer. Our integrated *in silico* and *in vitro* results confirmed the dual potential effect of the Montelukast both on virus entry into the host cell (Spike/ACE2) and on the main protease enzyme inhibition. The virus neutralization assay results showed that while no cytotoxicity of the Montelukast was observed at 12 μM concentration, the cell index time 50 (CIT_50_) value was delayed for 12 hours. Moreover, it was also shown that Favipiravir, a well-known antiviral used in COVID-19 therapy, should be used by 16-fold higher concentrations than Montelukast in order to have the same effect of Montelukast. The rapid use of new small molecules in the pandemic is very important today. Montelukast, whose pharmacokinetic and pharmacodynamic properties are very well characterized and has been widely used in the treatment of asthma since 1998, should urgently be completed in clinical phase studies and if its effect is proven in clinical phase studies, it should be used against COVID-19.

## 1. Introduction

The 2019 new coronavirus (SARS-CoV-2), was first reported in December 2019 in Wuhan (Hubei, China). It has quickly spread to other countries all around the world and effected more than 67 million people worldwide becoming an urgent global pandemic. Coronaviruses are enveloped, non-segmented positive-sense RNA viruses belonging to the family of Coronaviridae, the largest family in Nidovirales and widely distributed in humans, other mammals and birds, causing respiratory, enteric, hepatic and neurological diseases. Seven species of coronavirus are known to cause disease in humans. Four of them (229E, OC43, NL63, and HKU1) are common and they mostly cause common cold symptoms in immunocompetent individuals while the other three, SARS-CoV, MERS-CoV, and SARSCoV-2 cause serious symptoms and death.^1^ In addition to the common cold symptoms, SARS-COV-2 shows many clinical signs including severe pneumonia, clot formation RNAaemia and the incidence of endothelitis, fatigue, neurological and cardiac consequences.^2^

All coronaviruses have specific genes in ORF1 downstreams that encode proteins for viral replication, nucleocapsid and spikes development.^3^ SARS-CoV-2 has four structural proteins which are nucleocapsid, envelope, membrane and spike. These four proteins play a vital role during the viral infection.^4^ The Spike glycoprotein (S protein) located on the external surface of coronaviruses are responsible for the connection and entry of the virus to host cells.^1^ The S protein mediates receptor recognition, cell attachment, and fusion during viral infection. While the virus is in its natural environment, S protein of coronavirus is inactive. During viral infection, target cell proteases activate the S protein by cleaving it into S1 and S2 subunits, which are required to activate the membrane fusion domain after viral entry into target cells.^5^ The S1 subunit includes the receptor binding domain (RBD). This domain binds directly to the peptidase domain angiotensin converting enzyme 2 (ACE-2). S2 functions during membrane fusion.

The chymotrypsin-like cysteine protease called 3C-like protease (3CLpro) *aka* main protease (Mpro) in SARS-CoV-2 is a vital enzyme involved in processes such as the processing, assembly, and replication of the virus. Thus, Mpro is one of the ideal targets for drug design and development studies against SARS-CoV-2.^6^

One of the key characteristics of severe COVID-19 is increased cytokine production. It is thought that the severity of the disease is primarily associated with the cytokine storm, which is an aggressive immune response to the virus.^7^ The number of white blood cells, neutrophils, and levels of procalcitonin, C-reactive protein and other inflammatory indices like IL2, IL7, IL10, granulocyte-colony stimulating factor (GSCF), interferon inducible protein −10 (IP10), monocyte chemotactic protein-1 (MCP1), macrophage inflammatory protein-1α (MIP1A), and TNF are significantly higher in severe cases in patients with COVID-19.^8,9^ Specifically, IL-1β, IL-6, and IL-10 are the three most elevated cytokines in serious cases.^10,11^ One result of the cytokine storm is lung injury that can develop into acute lung injury or its more severe type (acute respiratory distress syndrome, ARDS).

Studies have shown the relation between COVID-19 and the most common chronic conditions such as diabetes, cardiovascular diseases, respiratory system diseases, immune system disorders, etc.^7,12^ Asthma and chronic obstructive pulmonary disease (COPD) are among the diseases of the respiratory system that are most emphasized. Asthma is a chronic inflammatory airway condition. There is significant evidence that represents the relation of asthmatic patients in the population with viral infections like rhinoviruses.^13–15^ Virus infections cause upper respiratory tract infection, like influenza A, rhinovirus, and respiratory syncytial virus (RSV) elevate local leukotriene levels.^16^ Leukotrienes, which play a role in the contraction of bronchial muscles, are effective in initiating and amplifying many biological responses, including mast cell cytokine secretion, macrophage activation, and dendritic cell maturation and migration. Leukotrienes (LTC4, LTD4 and LTE4), activated basophils, eosinophils, macrophages, and products of mast cells are types of lipids conjugated with peptides.^17^

LTD4 receptors belong to G protein-coupled receptor (GPCR) family. Montelukast is a selective leukotriene (D4) receptor antagonist which is a member of quinolines and it was approved by FDA as an oral tablet in 1998. It is a licensed drug used for allergic rhinitis, exercise-induced bronchospasm and especially prophylaxis and chronic treatment of asthma. As a result of LTD4 blockage, NF-κB pathway activation and release of the proinflammatory mediators (i.e., IL-6,8 and 10, TNF-a and MCP-1) decrease^3^. Considering these anti-inflammatory effects by leukotriene receptor inhibition and possible antiviral effects, Montelukast may be considered for the effective medication against SARS CoV-2.^18,19^ Some studies claim that Montelukast may play an immunomodulatory role as a leukotriene receptor inhibitor in treatment since one of the pathophysiological steps of severe COVID-19 cases is the cytokine storm resulting from excessive proinflammatory mediator releasing.^20,21^

Nowadays the concept of drug repurposing is an evolving technique in which approved drugs are commonly used to identify potential candidates for different diseases. Developing new drugs from scratch is a long process and thus impractical to cope with the current global challenge.^22^ Many drugs have several protein targets, and many diseases share molecular mechanisms that overlap each other. In this scenario, reusing drugs for new purposes and discovering their new uses by using computational approaches will dramatically lower the cost, time and risks of the drug development processes.^23^

Here, initially we explored the potential role of Montelukast in the management of SARS-CoV-2 infection with multiscale molecular modeling approaches and its promising results both in main protease and Spike/ACE2 interface encouraged us to perform further detailed *in vitro* experiments. The results of FRET-based biochemical assays, surface plasmon resonance (SPR), pseudovirus neutralization and virus neutralization experiments demonstrated the effect of Montelukast on SARS-CoV-2.

## 2. Results and Discussion

### 2.1 *In Silico* Drug Repurposing studies suggest usage of Montelukast against SARS-CoV-2

It has been recognized that the “single target-one molecule” approach is not very effective in treating complex diseases and alternative combination drugs are not appreciated due to toxicity and / or unwanted drug-drug interactions.^24,40^ The promising approach to these complex diseases is to develop single-multitarget compounds that a molecule may interact with multiple related selected target proteins simultaneously. As new drugs are expensive and time consuming to develop, repositioning / reusing drugs has emerged as an alternative approach. Thus, in our recent study^24^ we have screened FDA approved drugs and compounds in clinical investigation to identify effective single-multi-target molecules against HIV-1. For this aim, two important and essential proteins in the HIV-1 life cycle, CCR5 co-receptor (belongs to GPCR family) and HIV-1 protease (PR) enzyme, were targeted and about 25 potential hit compounds were identified. Montelukast was among the molecules we suggested as dual HIV-1 inhibitor with *in silico* simulations and that we purchased and added to our molecule library to evaluate *in silico* results. When Montelukast achieved successful results against SARS-CoV-2 main protease and Spike/ACE-2 targets in the preliminary virtual screening studies of our small molecule library belonging to our laboratory, we decided to carry it to further experimental studies.

Both noncovalent and covalent docking approaches are performed in the SARS-CoV-2 main protease since recent structural biology studies show that majority of the co-crystallized compounds at the main protease construct bonded interactions. Top-docking poses of Montelukast at the main protease and Spike/ACE2 targets were then used in 1 μsec all-atom molecular dynamics (MD) simulations. Figure 1 shows representative structure of Montelukast at the SARS-CoV-2 main protease target obtained from saved trajectories throughout the MD simulations initiated from non-covalent docking. Crucial residues in ligand interaction were found as His41, Met49, Asn142, Met165, Glu166, Leu167, Pro168, Phe185, Gln189 and Ala191 (Figure 2). In Figure 3, surface representation of representative complex structure of Montelukast with main protease show crucial interactions. Corresponding interactions obtained from simulations initiated from covalent docking were Asn142, Gly143, Ser144, Cys145, His164, Glu166, and Gln189 (Figure 4).

**Figure 1.**
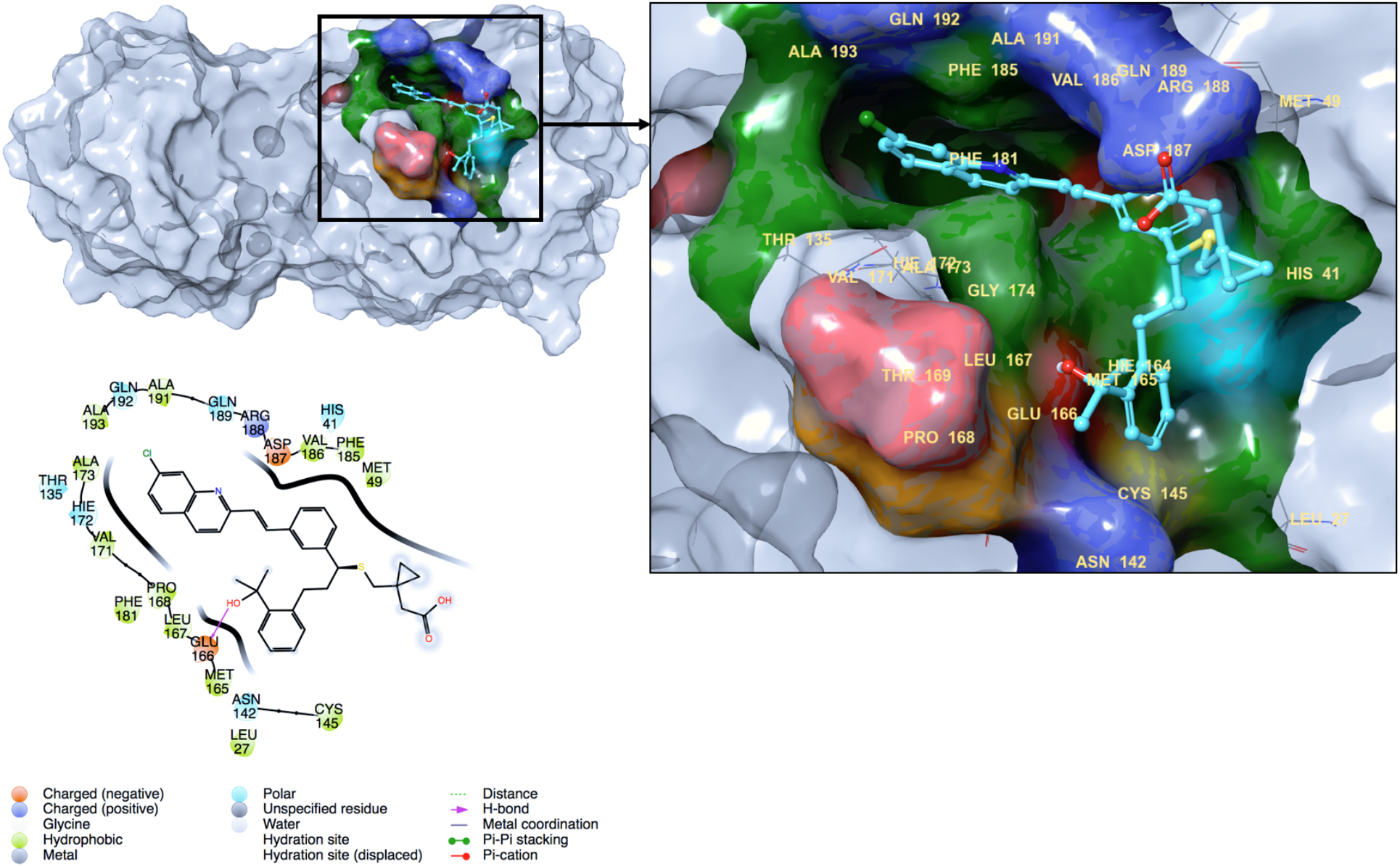
Representative complex structure of Montelukast at the binding pocket of the SARS-CoV-2 main protease obtained from saved trajectories of MD simulations initiated with its non-covalent top-docking pose. 2D ligand interaction diagram also represented.

**Figure 2.**
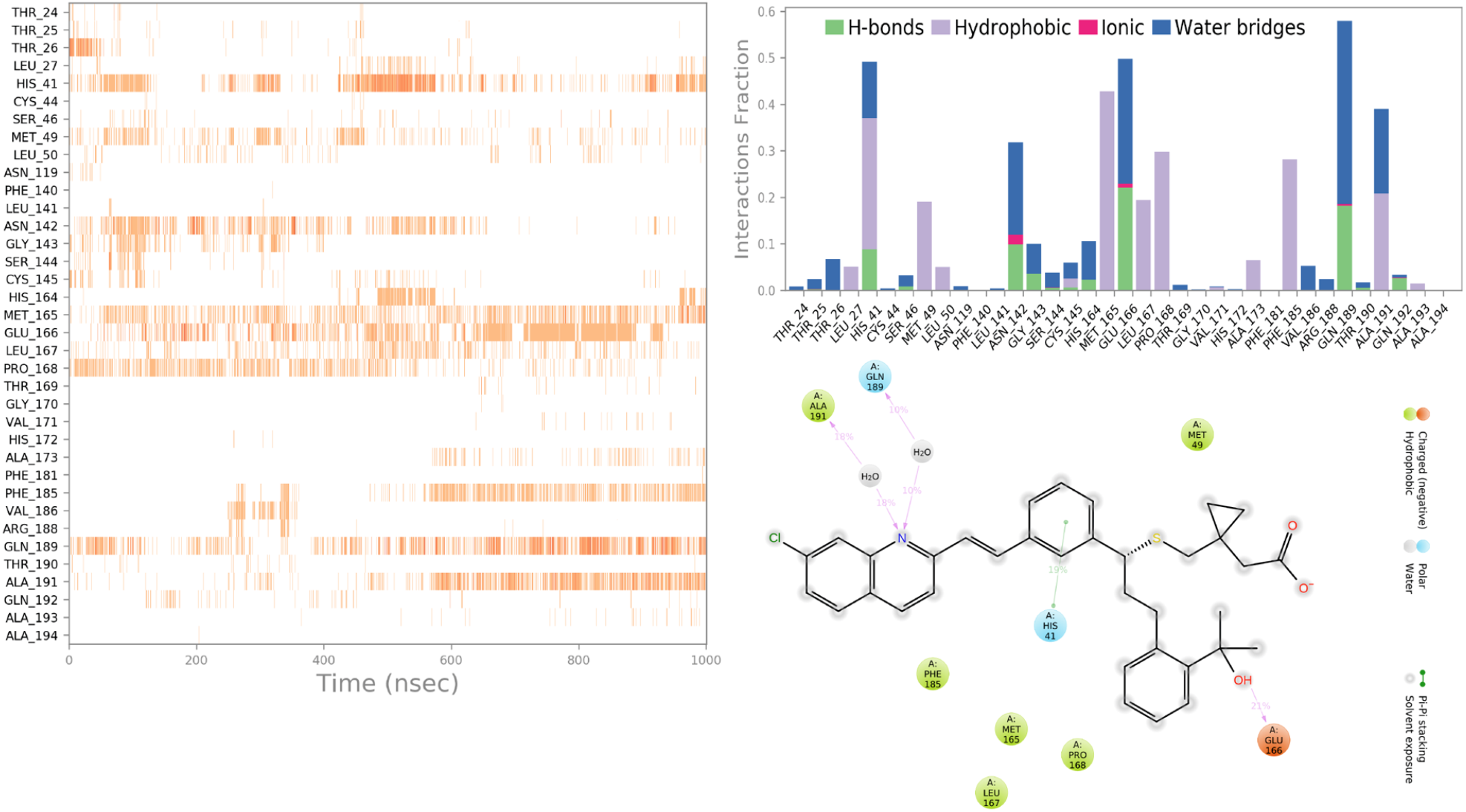
Crucial interactions of Montelukast during the MD simulations initiated with its non-covalent SARS-CoV-2 main protease top-docking pose. Protein interactions with the Montelukast is monitored during the simulation. The stacked bar charts are normalized over the course of the trajectory (i.e., a value of 0.5 represents that 50% of the simulation time the specific interaction is maintained).

**Figure 3.**
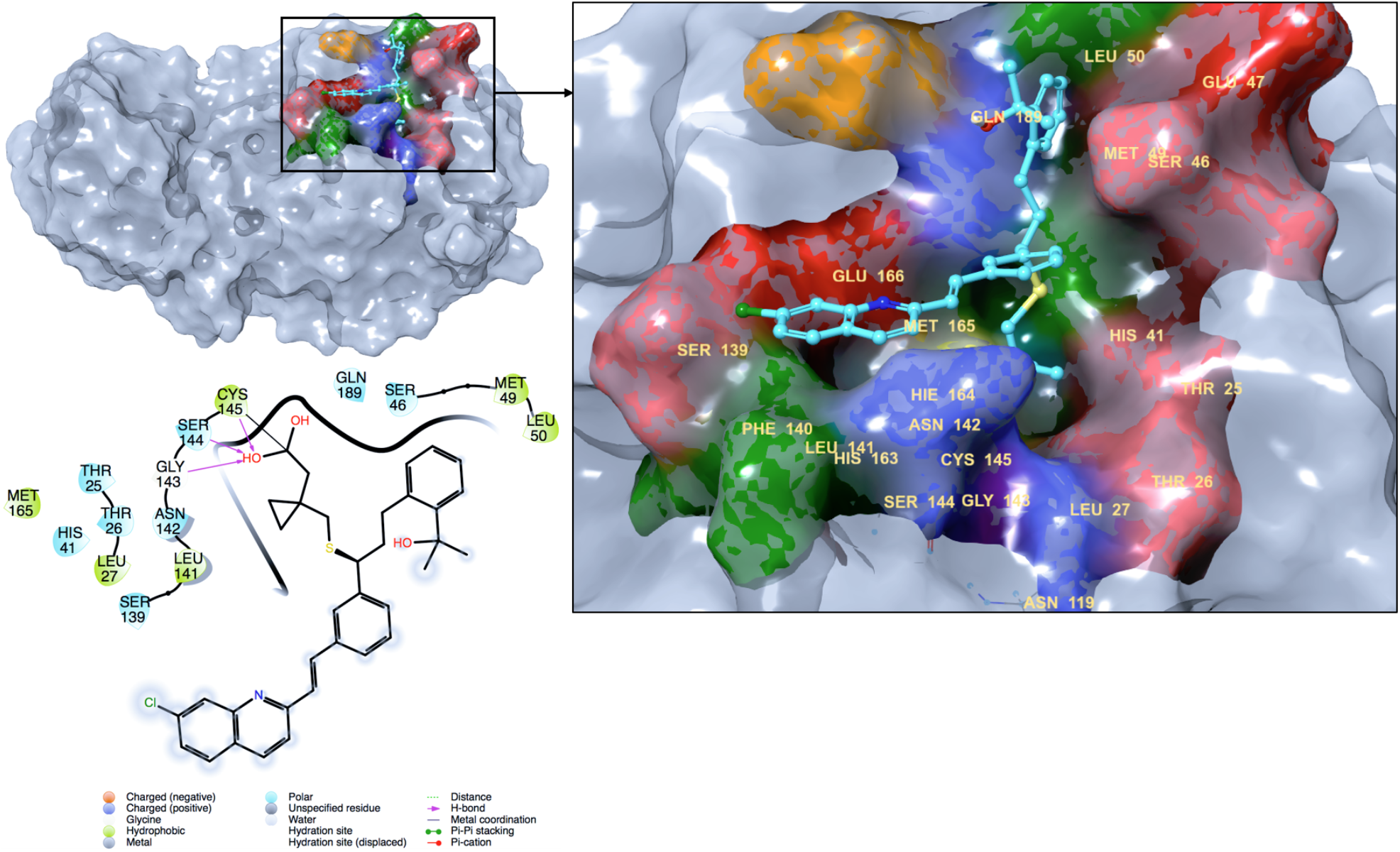
Representative complex structure of Montelukast at the binding pocket of the SARS-CoV-2 main protease obtained from saved trajectories of MD simulations initiated with its covalent top-docking pose. 2D ligand interaction diagram also represented.

**Figure 4.**
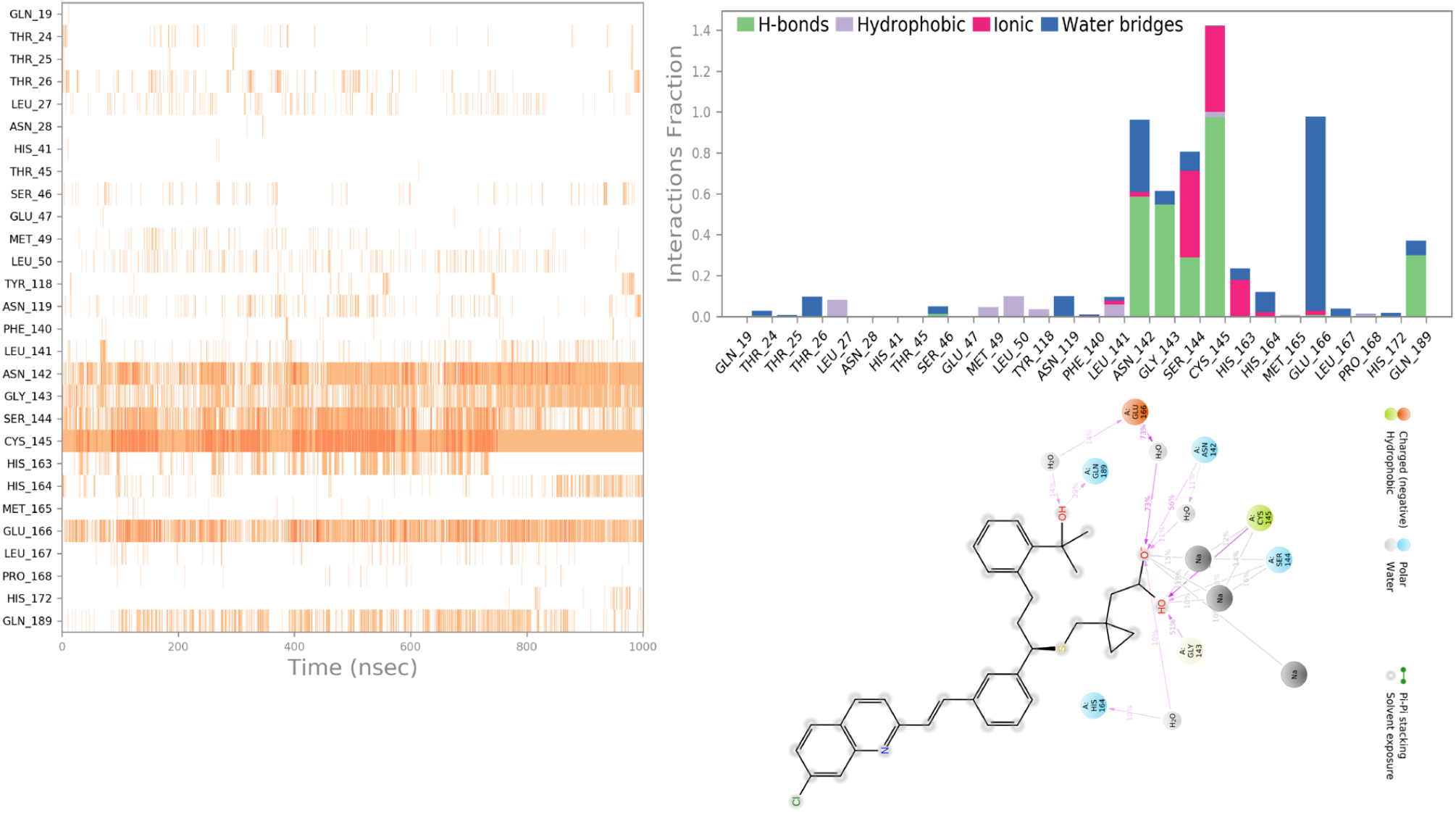
Crucial interactions of Montelukast during the MD simulations initiated with its covalent SARS-CoV-2 main protease top-docking pose. Protein interactions with the Montelukast is monitored during the simulation. The stacked bar charts are normalized over the course of the trajectory (i.e., a value of 0.5 represents that 50% of the simulation time the specific interaction is maintained).

The effect of Montelukast on the SARS-CoV-2 Spike/ACE-2 region is also investigated. Top-docking pose of Montelukast at the Spike/ACE-2 is used at the all-atom MD simulations. Figures 5 and 6 represent 3D and 2D ligand interaction diagrams. The important residues in ligand interactions were found as Lys26, Asp30, Val93, Pro389, Arg408, Lys417, Phe555, Asn556, and Arg559.

**Figure 5.**
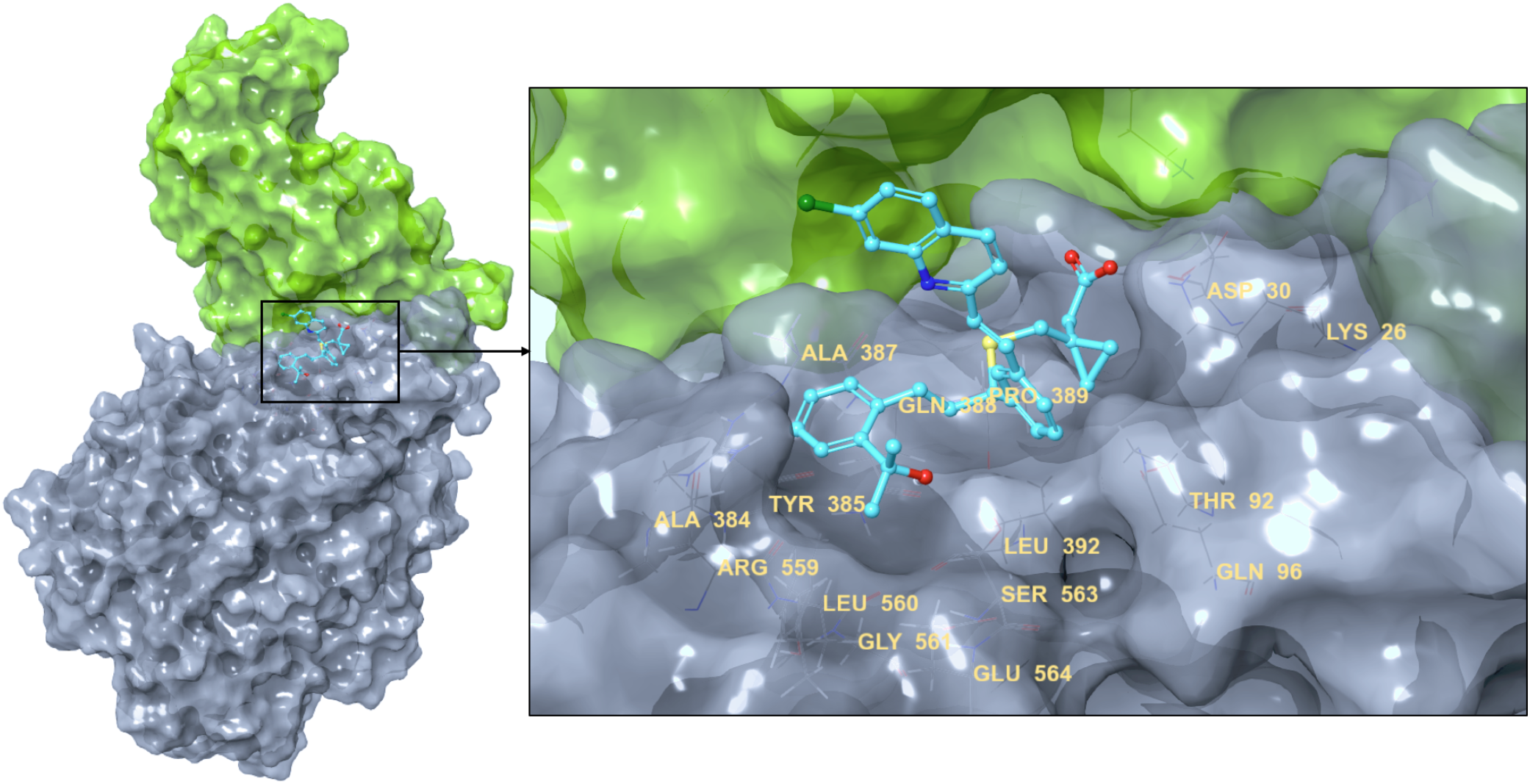
Representative complex structure of Montelukast at the SARS-CoV-2 Spike/ACE-2 interface obtained from saved trajectories of MD simulations initiated with its non-covalent top-docking pose.

**Figure 6.**
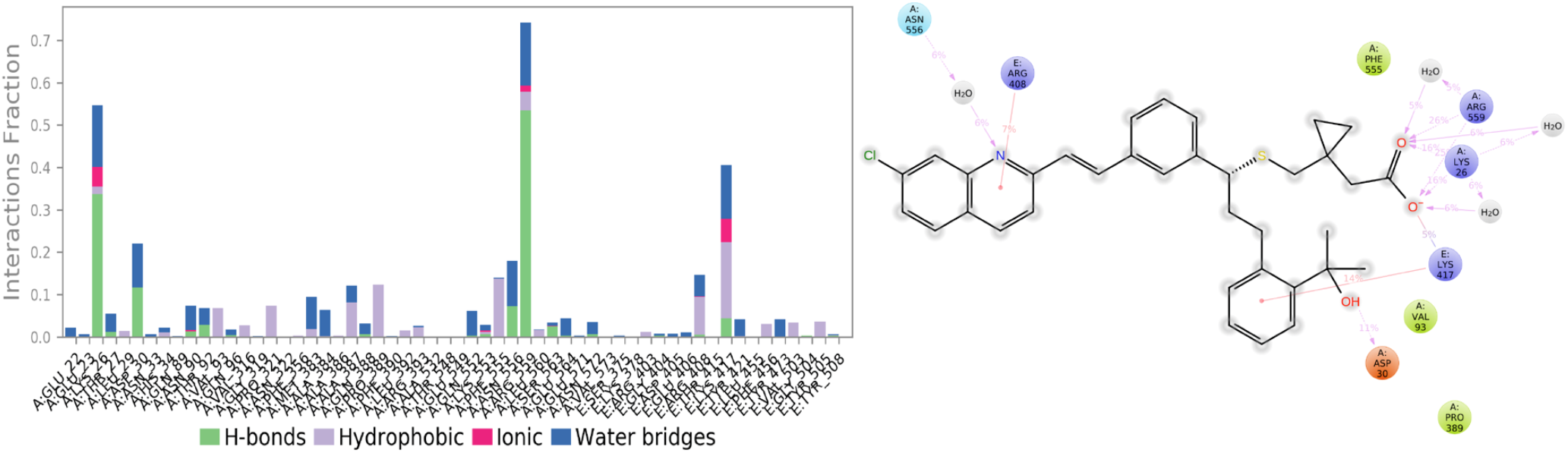
Crucial interactions of Montelukast during the MD simulations initiated with its SARS-CoV-2 Spike/ACE-2 top-docking pose. The stacked bar charts are normalized over the course of the trajectory (i.e., a value of 0.5 represents that 50% of the simulation time the specific interaction is maintained).

The binding energies of Montelukast at the main protease and Spike/ACE2 targets were also investigated. Average MM/GBSA binding energy of Montelukast at the binding cavity of main protease was measured as −79.60 ±8.66 kcal/mol. When we increased the simulation time (i.e., simulations are repeated with 3-fold), 3 μsec simulations also showed that Montelukast maintains its interactions with the crucial residues at the binding pocket of Mpro. Average MM/GBSA binding energy was calculated as −84.86 ±9.22 kcal/mol. The corresponding average MM/GBSA value of Montelukast at the Spike/ACE-2 was calculated as −43.93±7.66 kcal/mol. (Figure 7)

**Figure 7.**
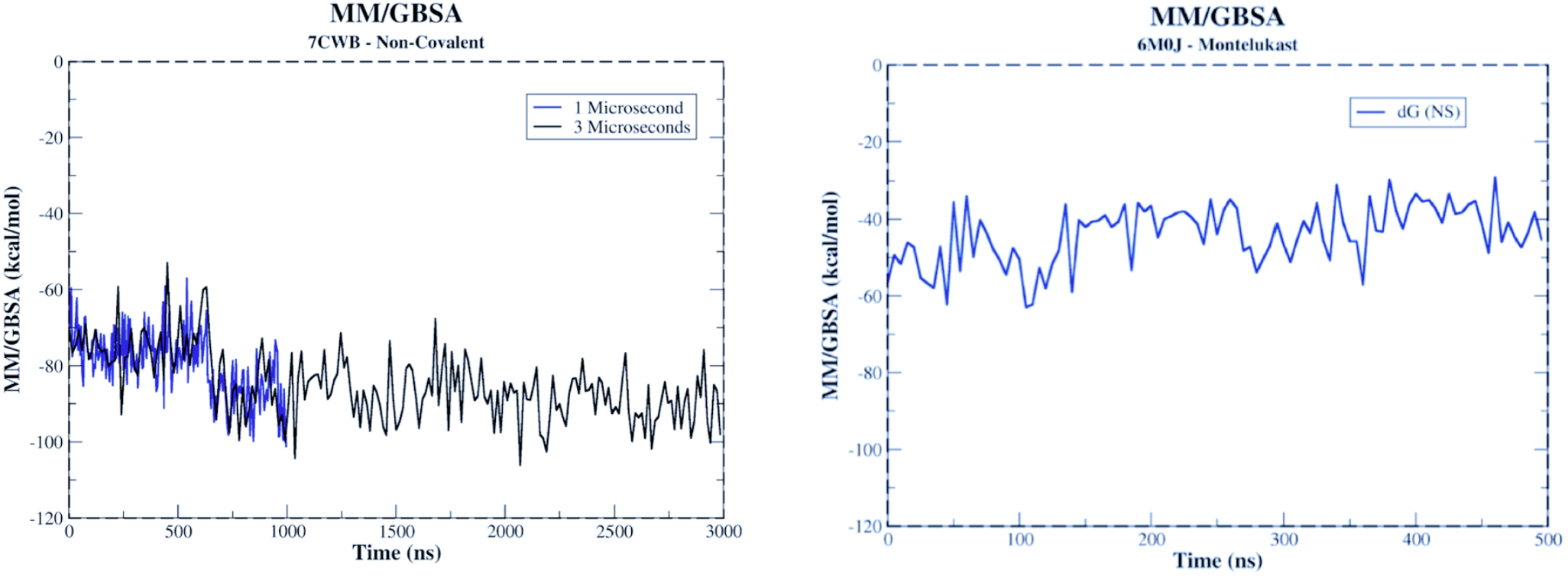
MM/GBSA analysis of Montelukast at the SARS-CoV-2 main protease (left) and Spike/ACE-2 (right) targets throughout the MD simulations. Mpro simulations are repeated by increasing the simulation time by 3-folds. Results show that Montelukast maintains its interactions during 3 μsec simulations.

### 2.2 Fluorescence Resonance Energy Transfer (FRET)-based main protease enzyme inhibition assay shows 74% loss of enzyme activity when using 100 μM Montelukast

The main protease assay revealed that the Montelukast molecule inhibits the main protease enzyme activity in concentration dependent manner. The maximum inhibitory concentration was 100 μM and inhibited the enzyme activity by 74.04%. At 50 μM concentration inhibition the enzyme activity inhibition was 60.59%. (Figure 8) GC376 is a broad-spectrum antiviral which is used as a positive control molecule at the used assay and it showed 88.59% enzyme activity inhibition at 100 μM. Nonlinear regression analysis of five different concentrations revealed that 50% inhibition concentration (IC_50_) of Montelukast molecule is 28.36 μM (Figure 8). Experiments are repeated by 6-times in IC_50_ measurement.

**Figure 8.**
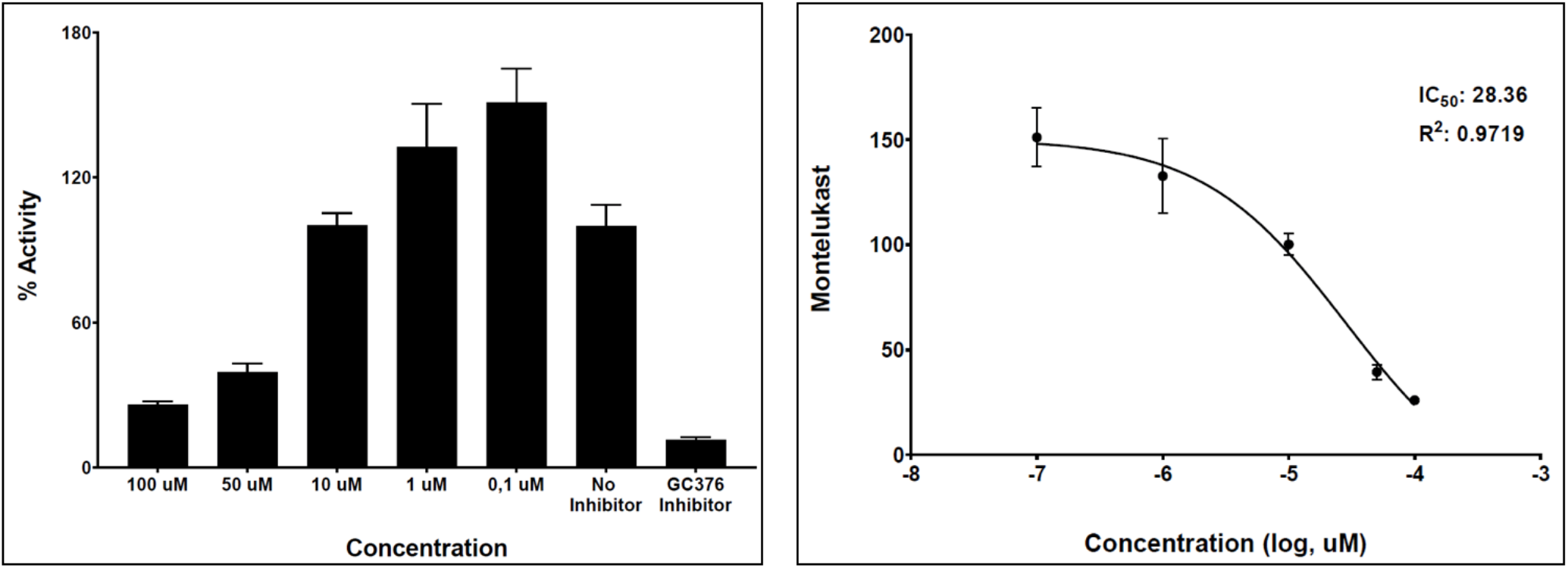
**(left)** 3CL Protease activity in the presence of Montelukast with ranging concentrations. “Inhibitory Activity” shows the inhibited 3CL (main protease) enzyme activity percentage. “No inhibitor” represents the 3CL protease activity without any inhibitors, GC376 inhibitor is a broadspectrum antiviral used for comparison. **(right)** Dose-response curve of Montelukast against 3CL protease. Experiments are repeated at least 3 times.

### 2.3 The surface plasmon resonance (SPR) assays demonstrate Montelukast binding to SARS-CoV-2 main protease

In the present study, together with FRET-based binding analyses of Montelukast at the SARS-CoV-2 main protease, surface plasmon resonance (SPR) spectroscopy was also used to evaluate the binding kinetics and affinity of this interaction. Biosensor technology from SPR has become an important tool for drug design and discovery. SPR techniques are used for a broad range of applications including assessing the binding kinetics and affinity of an interaction, specificity tests, ligand screening, as well as analyte active binding concentration measurements. It can be used for the aim of drug screening for several diseases including COVID-19. Here, SPR was used to estimate the potential role of Montelukast in the management of SARS-CoV-2 infection and its binding kinetics on main protease after analysis of multiscale molecular modeling studies and FRET-based main protease enzyme inhibition assays.

Solvent correction for 9.2% DMSO was shown in Figure 9. The affinity of Montelukast to immobilized Mpro was determined using a 1:1 steady-state binding affinity interaction model. A concentration series ranging from 900 μM to 11 μM (3-fold dilution) was injected over immobilized Mpro for 60 sec followed by a 120 sec dissociation phase. The responses obtained from each Montelukast concentration were plotted against concentration using the Biacore T200 evaluation software and was evaluated using a 1:1 steady-state binding model. Montelukast was identified as a specific binder to main protease (Figure 9). Its K_D_ value was measured as 23.5 μM which fits well with the FRET-based determined IC_50_ value. Squared-shape of sensorgrams shows that both Montelukast binding to MPro and complex dissociation are fast processes. This kind of binding behaviour is, however, relatively common for small molecules.

**Figure 9.**
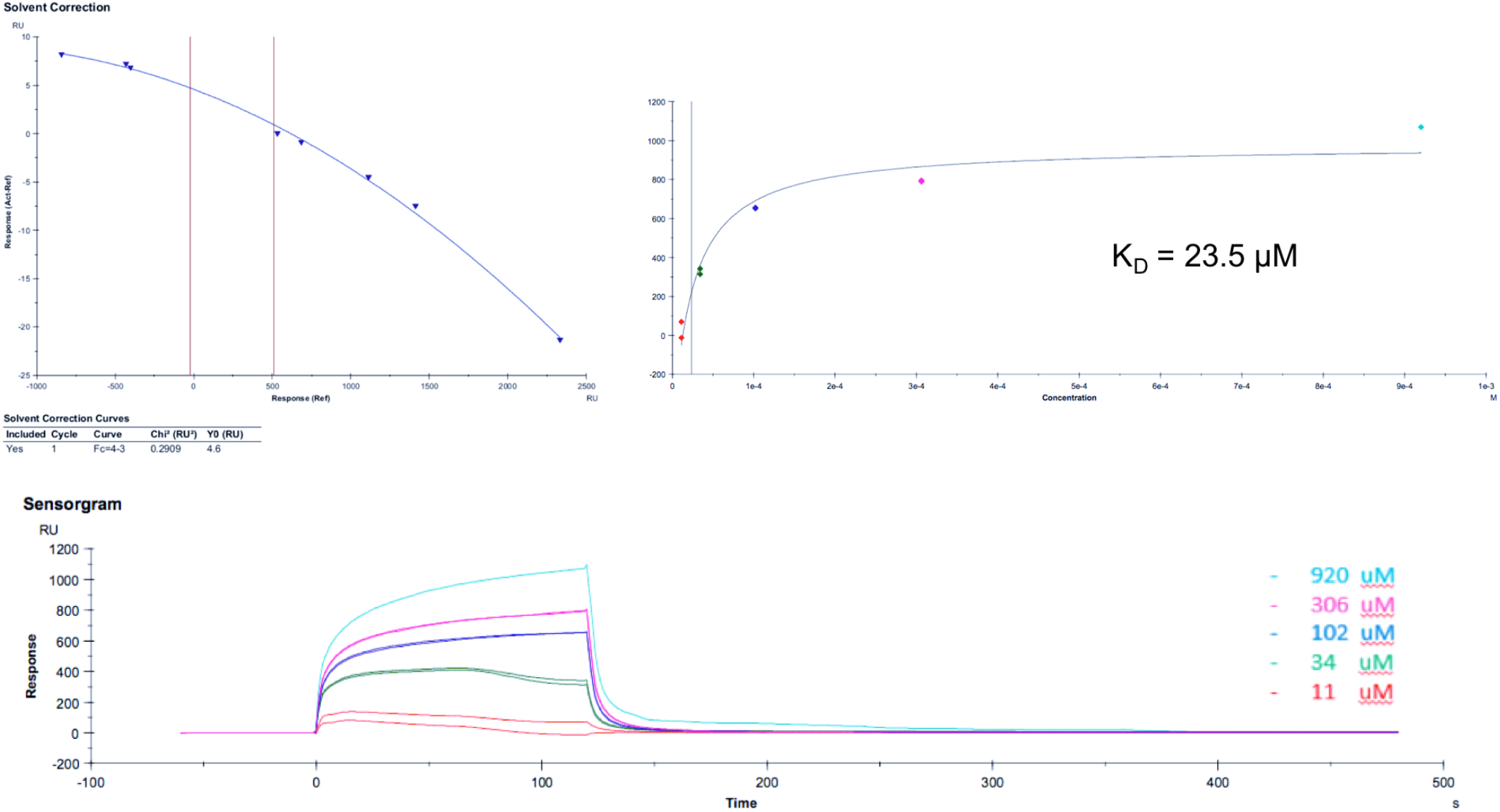
**(top-left)** Solvent correction for dimethyl sulfoxide (DMSO) for the percentage of DMSO between 10% and 8.6%. The sample was dissolved in 9.2% DMSO. **(top-right)** 1:1 steady-state binding model of Montelukast at the 3C-like protease, **(bottom)** Subtracted and correction sensograms of Montelukast binding curves for 3C-like protease.

The observed concentration-dependent binding responses, from the preliminary results, indicate that Montelukast molecule interacts with main protease with an affinity in the micro-molar range. According to the sensorgrams, the interactions do not reach a plateau (equilibrium phase) and also the small decrease of the sensorgrams at the end of the binding phase indicate that some aggregation issue might be present. Therefore, the determination of the exact binding affinity constant of the Montelukast to main protease is restrained.

### 2.4. Neutralization potential of Montelukast against SARS-CoV-2 was confirmed by pseudovirus neutralization on HEK293T/hACE2

#### Cell Viability

To investigate the dose-response relationship of Montelukast using the cell lines of HEK293T, Vero E6, Calu-3, and A549, the cell viability was measured upon 24, 48, and 72 hr exposure periods by MTT calorimetric assay. As shown in the Figure 10, HEK293T cell monolayer was found to be the most sensitive cell line across the Montelukast exposure and followed by the Vero E6, A549 and Calu-3. After 24, 48, and 72 hrs exposure periods, the IC_50_ value of Montelukast toward the used cell lines were given in the Table 1. The result of the cell viability assay allows us to determine the concentration ranges for the pseudovirus neutralization assay.

**Figure 10.**
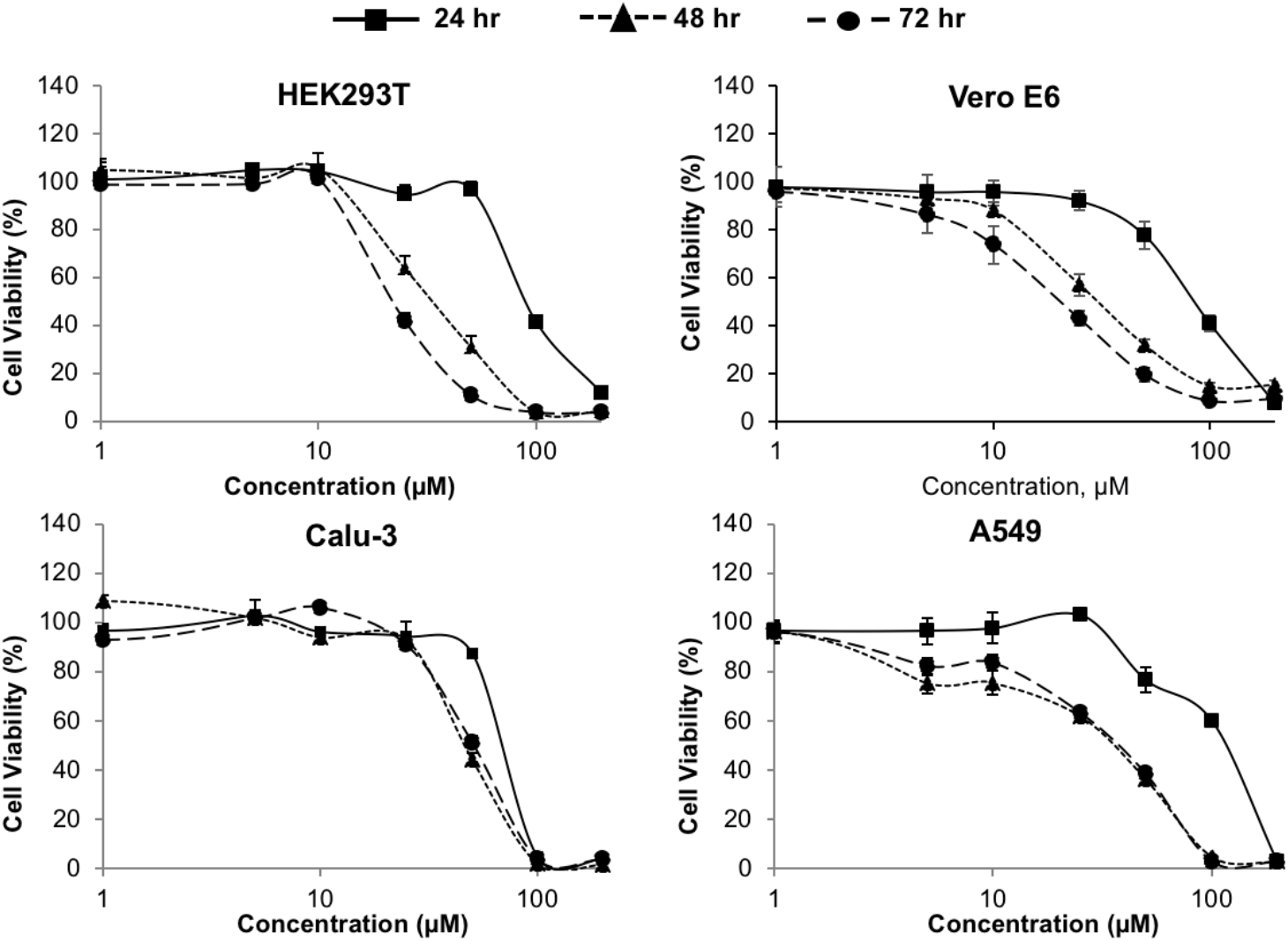
Cell viability of HEK293T, Vero E6, Calu-3, and A549 upon exposure to Montelukast at the concentration range of 1 to 200 μM for 24, 48, and 72 hrs. It was measured by MTT colorimetric assay. Each dose was tested in triplicate and error bars indicate SEM of triplicates.

**Table 1.** The IC_50_ values of Montelukast toward the used HEK293T, Vero E6, Calu-3, and A549 cells lines obtained from dose-response curves. Mean ± SD values were calculated from three independent experiments carried out triplicate.

| Exposure, hr | IC<sub>50</sub> values of Montelukast (μM) | | | |
| --- | --- | --- | --- | --- |
|  | HEK293T | Vero E6 | Calu-3 | A549 |
| **24** | 92.2 ± 0.8 | 92.39 ± 3.8 | 72.7 ± 0.3 | 117.6 ± 1.4 |
| **48** | 35.3 ± 4.0 | 38.80 ± 2.5 | 47.1 ± 1.3 | 38.2 ± 1.9 |
| **72** | 23.0 ± 0.8 | 28.11 ± 3.1 | 50.7 ± 1.9 | 38.5 ± 0.8 |

##### Pseudovirus neutralization

Pseudovirus neutralization on HEK293T/hACE2 was performed using Montelukast at the concentration ranges of 1 to 50 μM (Figure 11). Inhibition of entry of HEK293T/hACE2 cells were pretreated with Montelukast and transduced with SARS-CoV-2 S pseudovirion. To evaluate the cell viability at the same concentrations which were used for neutralization, the luciferase activity was measured 72 hr post transduction by CellTiter-Glo Luminescent Cell Viability Assay Kit (Figure 11A). The entry efficiency of SARS-CoV-2 pseudoviruses without any treatment was taken as 100%. The representative plug formations at different concentrations was shown in the photos which were taken with magnification 10X during neutralization period (Figure 11B). The 50% effective neutralization concentration of Montelukast was found as 54.04 μM, therefore the neutralization potential of montelukast against SARS-CoV-2 was confirmed.

**Figure 11.**
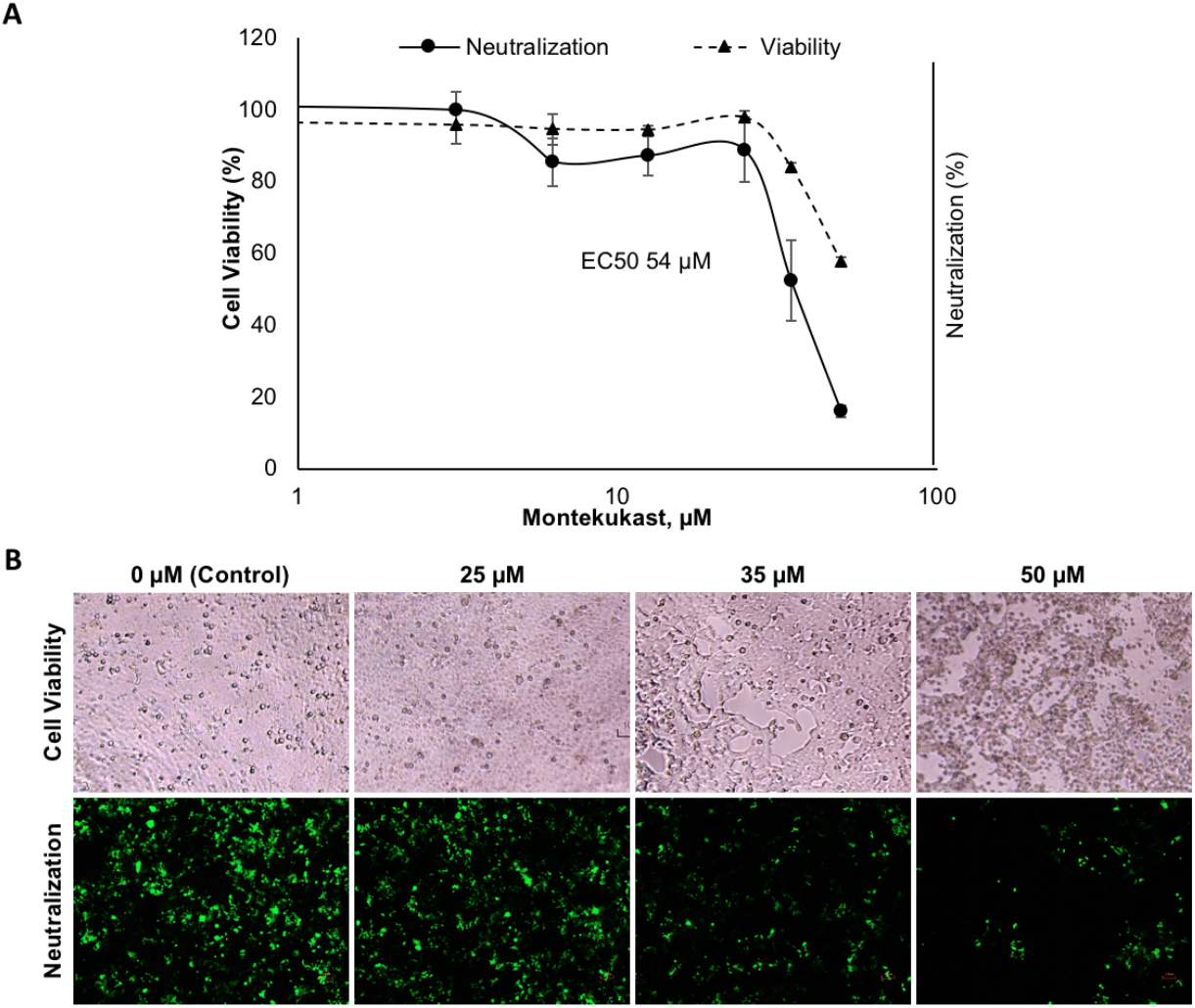
Pseudovirus neutralization on HEK293T/hACE2 by Montelukast. **(A)** Inhibition of entry of HEK293T/hACE2 cells were pretreated with different concentrations of Montelukast and transduced with SARS-CoV-2 S pseudovirion. The luciferase activity was measured 72 h post transduction. The entry efficiency of SARS-CoV-2 pseudoviruses without any treatment was taken as 100%. Each dose was tested in triplicate and error bars indicate SEM of triplicates. (**B)** The representative images for the cell viability and neutralization were shown upon neutralization period, 72 hr. Magnification 10X.

Pseudoviruses are useful tools due to especially for emerging and re-emerging viruses, their safety and versatility. To increase the transfection and infection potency in the development stage of the pseudovirus, the main factors including selection of plasmids, cell types, cell numbers, virus inoculum needs to be optimized. In this study, pseudovirus neutralization assay was developed for screening computationally selected drug, Montelukast, as potent inhibitor of SARS-CoV-2 main protease.

### 2.5. Virus Neutralization Assay Using xCELLigence MP Real Time Cell Analyzer Demonstrates the Effective Virus Neutralization Concentration of Montelukast

Neutralization assay was performed based on impedance using xCELLigence MP real time cell analyzer equipment (RTCA). VERO E6 cells were used. Neutralization assay was performed based on impedance (resistance to alternating current) using xCELLigence MP RTCA. The impedance is expressed as arbitrary units called cell index (CI). Real-time cell analysis result of Montelukast without virus incubation during the 150 hours shows that Montelukast can be used up to 23 μM without cytotoxic effects (Figure 12) Figure 13 shows real-time cell analysis result of Montelukast with the SARS-CoV-2. Data was collected for 150 hours with intervals of 15 min.

**Figure 12.**
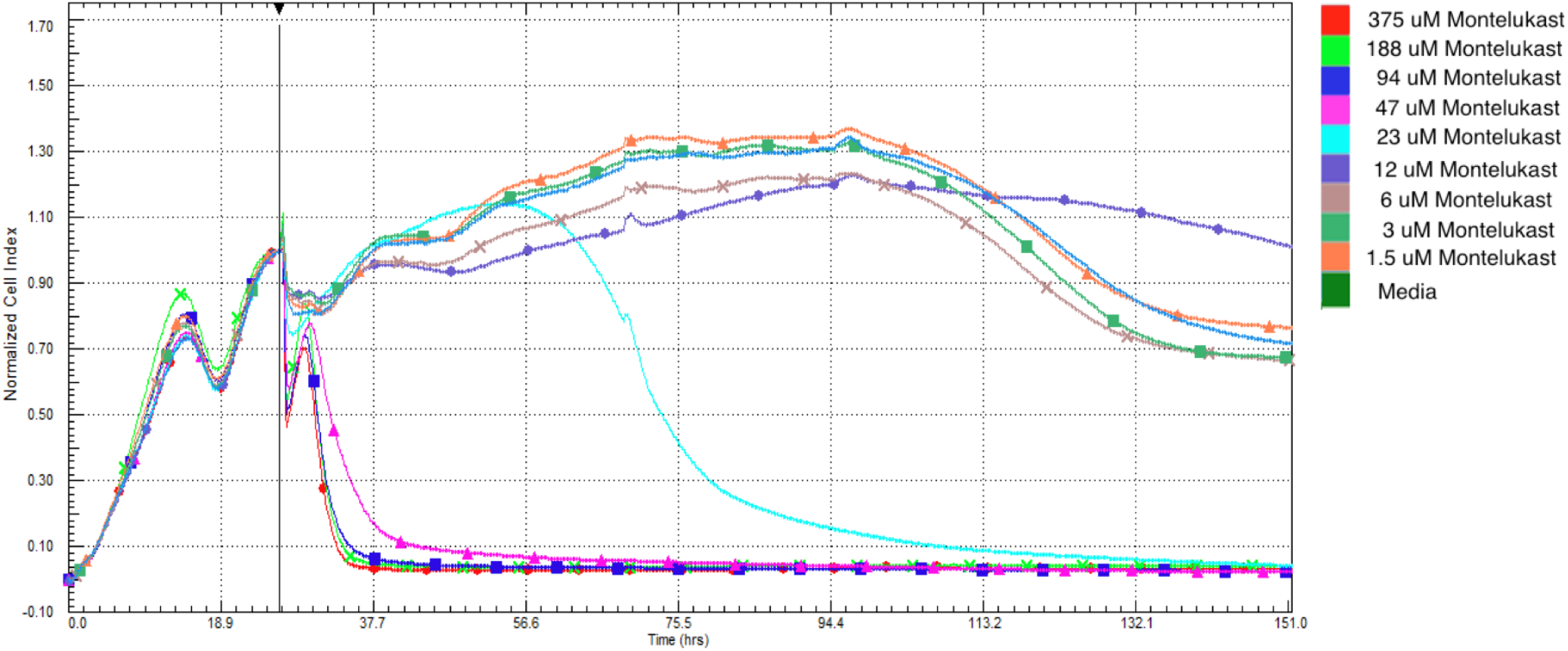
Real-time cell analysis result of Montelukast without virus incubation. Data was collected for 150 hours with intervals of 15 min. At the end of the period, the experiment was terminated and the data obtained were analyzed using RTCA Software Pro software. The samples in the figure are indicated according to their color codes.

**Figure 13.**
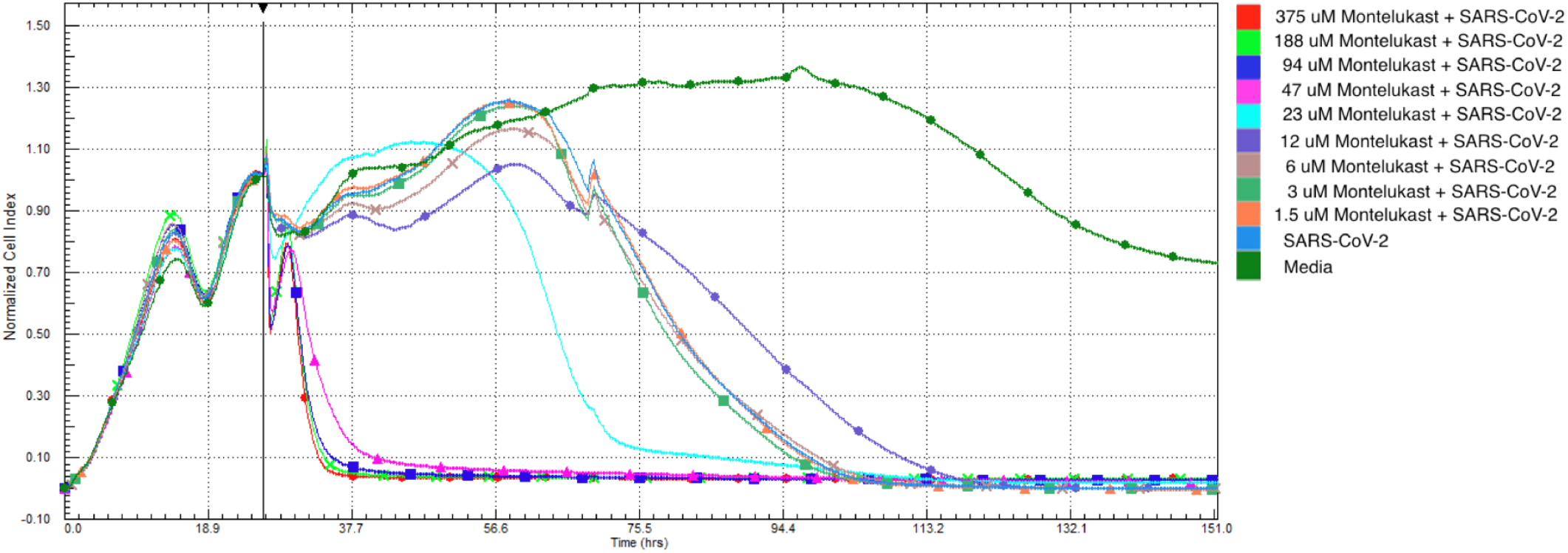
Real-time cell analysis result of Montelukast. Data was collected for 150 hours with intervals of 15 min. At the end of the period, the experiment was terminated and the data obtained were analyzed using RTCA Software Pro software. The samples in the figure are indicated according to their color codes.

Results show that compound demonstrates neutralization effect at 12 μM. In this concentration, it is not cytotoxic. The cell index time 50 (CIT_50_) is the time required for the CI to decrease by 50% after virus infection. While no cytotoxicity of the molecule was observed at 12 μM concentration, the CIT_50_ value was delayed for 12 hours. (Figure 14) When the same experiment is carried out for the Favipiravir with same conditions, which is the one of the mostly used antiviral at the COVID-19, we observed that Favipiravir should be used in 16-fold more higher doses than Montelukast in order to show the same effect (i.e., around 12 hours delay in CIT_50_), (Figures 15 and 16).

**Figure 14.**
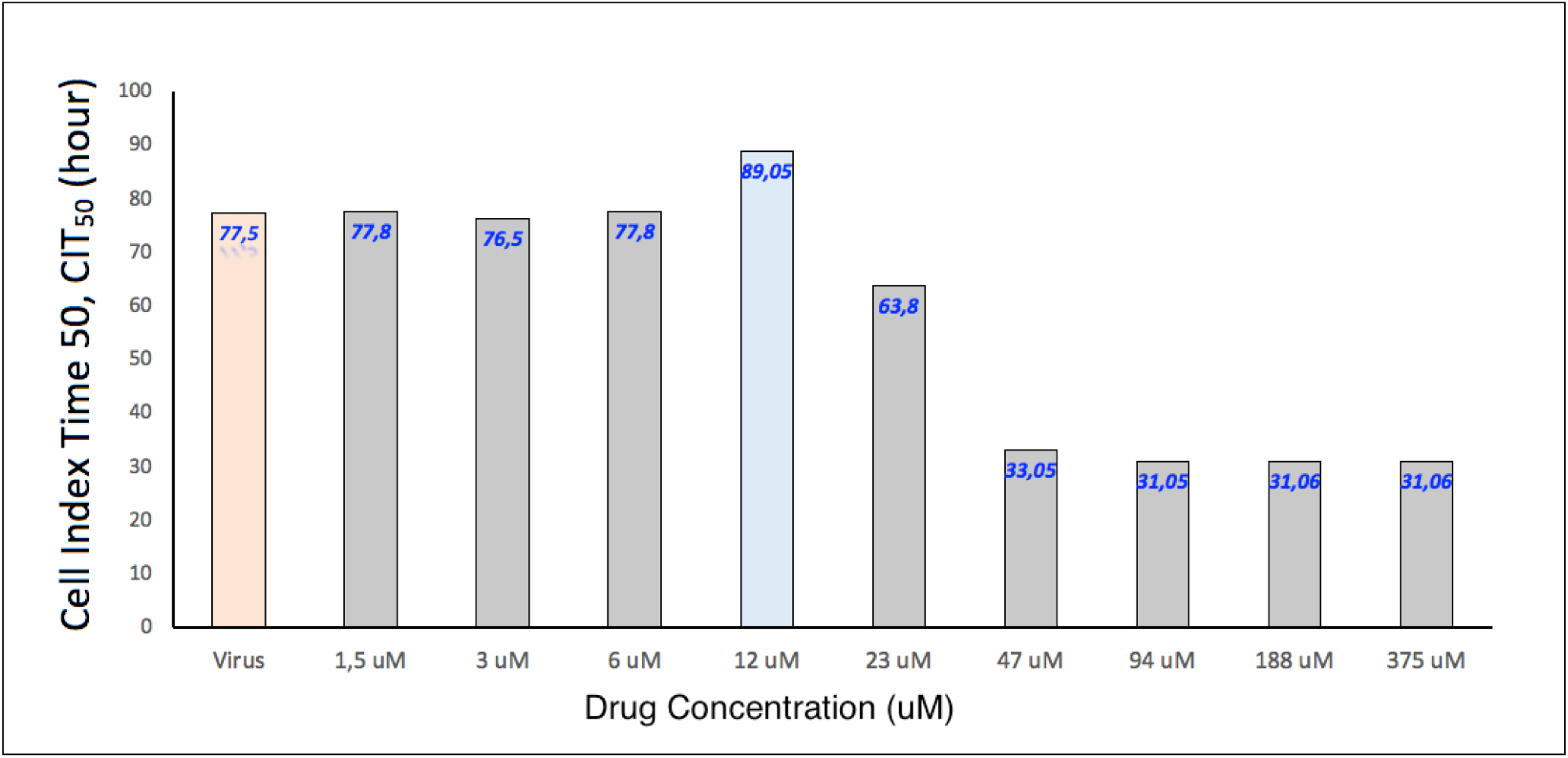
Comparison of the CIT_50_ values of the SARS-CoV-2 virus neutralization efficacy study obtained with RTCA of different concentrations of the Montelukast molecule.

**Figure 15.**
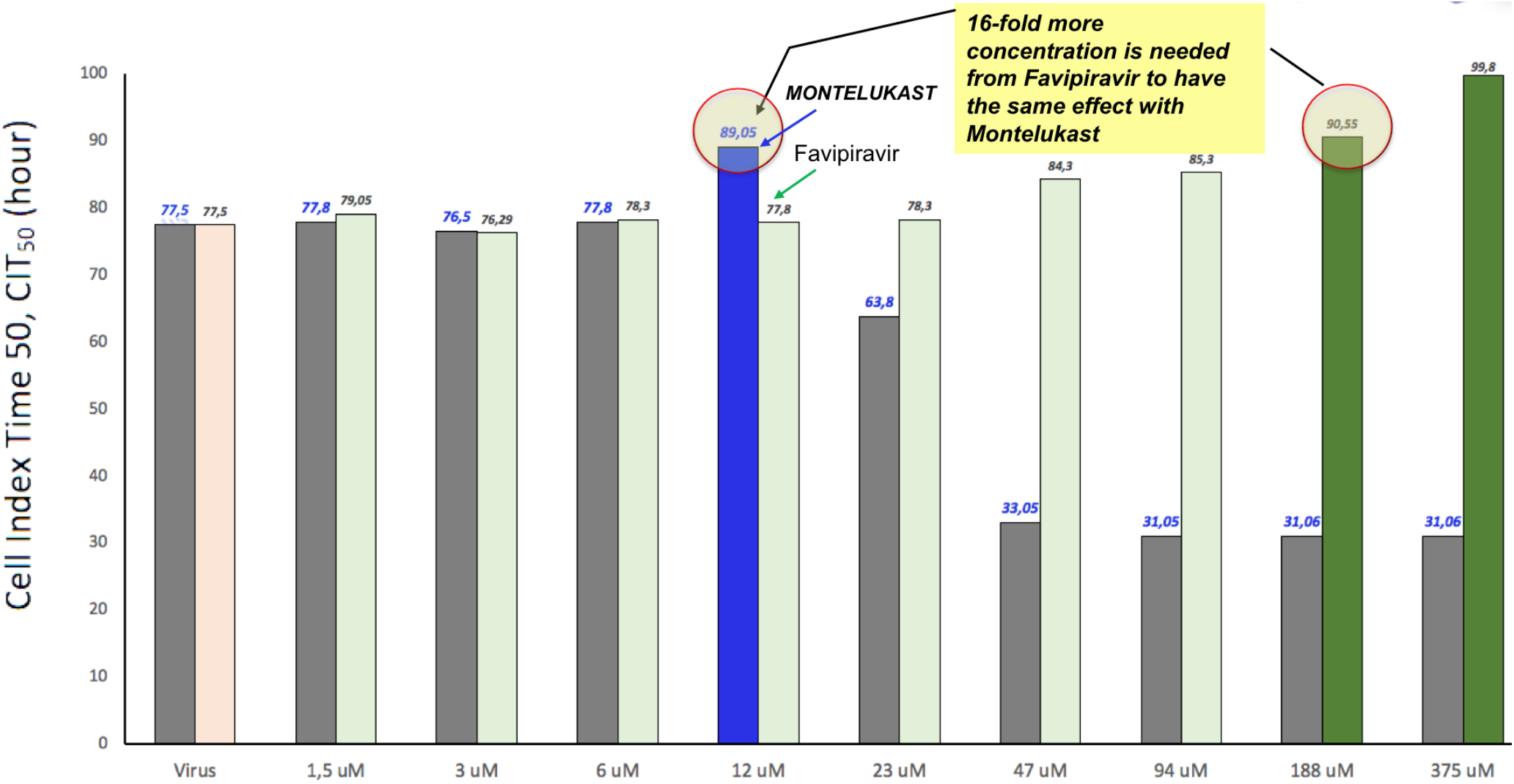
Comparison of the CIT_50_ values of the SARS-CoV-2 virus neutralization efficacy study obtained with RTCA of different concentrations of the Montelukast and Favipiravir molecule. While gray bars show Montelukast, pale green colored bars represent Favipiravir. Effective concentrations for Montelukast and Favipiravir are represented in different colors, dark blue (Montelukast), dark green (Favipiravir).

The investigation of anti-viral activity and pseudovirus and virus neutralization potential therapeutic agents against the live SARS-CoV-2 has to be performed under biosafety level 3 conditions because of its high pathogenicity and infectivity.^25,26^ Thus, in the current study, our integrated *in silico* and combined *in vitro* experiments show the effect of Montelukast at the SARS-CoV-2.

## 3. Methods

### 3.1. Molecular Modeling Studies

Before the molecular docking and MD simulations, both ligand structure and used target protein structures were prepared. Montelukast structure was downloaded from PubChem (https://pubchem.ncbi.nlm.nih.gov/) and LigPrep module of the Maestro molecular modeling package was used in ligand preparation with OPLS3e force field. Epik was used in the determination of the protonation states at neutral pH.^27^ While the crystal structure of main protease which is recently solved by our research group (PDB, 7CWB) in near physiological temperature ^28^ was used in docking and MD simulations, 6M0J coded structure was used in drug docking and all-atom MD simulations for Spike/ACE-2 region. Protein preparation tool of Maestro was used in both targets at physiological pH. Bond orders are assigned, and hydrogens were added. Disulfide bonds were created and missing side chains were fixed using Prime.^29^ Water molecules beyond 5 Å from hetero groups were removed. PROPKA was used in the protonation states of the residues. OPLS3e force field ^30^ was used in the restrain minimization with heavy atom convergence of 0.3 Å.

#### 3.1.1 Noncovalent docking

The prepared target proteins and ligand structure were used for molecular docking simulations. We performed a grid-based docking method (Glide/SP) at the docking.^31^ Binding site of the main protease was defined by centering grids at the centroid of a set of three crucial residues in ligand binding, namely His41, Cys145, and Glu166. Ali and Vijayan^32^ stated a very strong and sustained salt bridge interactions between Lys417 of SARS-CoV-2 Spike RBD and Asp30 of ACE-2. Thus, the corresponding residues at the Spike/ACE-2 were used in grid generation. Top-docking poses obtained from noncovalent docking are used in all-atom MD simulations.

#### 3.1.2. Covalent docking

When the ligand forms a covalent bond with a binding pocket residue, the binding energy of the ligand is not only from the construction of a covalent bond but also from stabilizing nonbonding interactions. Here, we used Covalent Docking module from Maestro which selects the top-covalent bond poses using Prime energy model.^33^ In this method, docking starts with Glide docking with the reactive residue trimmed to Alanine residue. The reactive residue of the target protein is then added and sampled to form a covalent bond with the ligand in different poses. Formed poses with covalent bond are minimized using the Prime VSGB2.0 energy model to score the top-covalent poses. In covalent docking, ligand binding site was detected from three crucial residues at the main protease namely His41, Cys145, and Glu166. As reaction type “nucleophilic addition to a double bond” option is selected with the guidance of the covalent bonded co-crystalized structures of inhibitors at the main protease. In docking process, default parameters were used.

#### 3.1.3 MD Simulations and MM/GBSA analyses

The selected docking poses of Montelukast at the main protease and Spike/ACE-2 targets were used in MD simulations. The used top-docking poses of complex structures were placed in simulation boxes with orthorhombic box (box size was calculated based on buffer distance of 10.0 Å) and solvated with TIP3P water models. The simulation systems were neutralized by the addition of counter ions and 0.15 M NaCl solution was used. Desmond program was used for all-atom MD simulations.^34^ Before the production run, the systems were equilibrated using the default relaxation protocol of the Desmond. Simulations were performed at constant physiological temperature (310 K) and constant pressure (1.01325 bar). For this aim, NPT ensemble was used with Nose–Hoover thermostat^35,36^ and Martyna-Tobias-Klein barostat^37^. Smooth particle mesh Ewald method^38^ was used to calculate long-range electrostatic interactions with periodic boundary conditions. The cutoff distance was set to 9.0 Å for short range electrostatics and Lennard-Jones interactions. The RESPA multi-step integrator was used. The time steps were varied for interaction types (bonded and near, 2 fs; far, 6 fs). Production time of the simulation was 1 μs for main protease and 0.5 μs for Spike/ACE-2 simulations. In the Spike/ACE2 target protein (PDB, 6M0J), glycans were included at the simulations. The OPLS3e force field was used in simulations. 2000 trajectory frames were recorded with equal intervals during the simulations. The average Molecular Mechanics Generalized Born Surface Area (MM/GBSA) binding free energy of the Montelukast were calculated for 200 trajectory frames throughout the simulations. VSGB 2.0 solvation model was utilized during MM/GBSA calculations.

### 3.2 The 3CL Protease enzyme inhibition assay

For enzyme inhibition assessment Fluorescence Resonance Energy Transfer (FRET)-based cleavage assay by using 3CL Protease assay Kit (#79955-1 and #79955-2, BPS Bioscience, San Diego CA, USA) was used. 100mM stock concentration of Montelukast was solved with dimethyl sulfoxide (DMSO), then diluted to working concentrations ranging 100nM to 100μM with 1X assay buffer (20 mM Tris, 100 mM NaCl, 1 mM EDTA, 1 mM DTT, pH 7.3) as manufacturer suggested. Final DMSO concentrations were below 1% for each tested concentration. 15μl of 3CL Protease enzyme was distributed to each well except “Blanks”. GC376 was used as an inhibitor control. 5μl GC376 (50μM) was added to the wells designated as inhibitor control. 5μl of inhibitor in different concentrations (100nM-10nM-1nM-1 μM-10μM-100μM) was added to their relative wells, and a 1X assay buffer/DMSO mixture was added to blanks and positive controls. 250μM 3CL protease substrate was added to each well to start the reaction, and its final concentration was 50μM in 25μl volume. After 4 hours of incubation at room temperature, fluorescence was measured by a microtiter plate-reader (Hidex Sense Multi Mode Reader, Finland) at a wavelength of 360nm for excitation and 460nm for emission. Blank values are subtracted from values of all other wells. Percentage inhibitory activity of each concentration were calculated, the fluorescence values from GC376 inhibitor control was set as zero percent activity and the fluorescence value from no inhibitor control was set as 100% activity. The IC_50_ (i.e., half-maximal inhibitory concentration) value was determined by also 3CL inhibitory screening assay. Absorbance values were recorded and corresponding IC_50_ value was calculated by dose response – inhibition curve and nonlinear regression analysis. The results were plotted with GraphPad Prism 8.0 software (GraphPad, San Diego CA, USA).

### 3.3 Surface Plasmon Resonance (SPR)

Biacore T200 spectrometer Cytiva (Uppsala, Sweden) instrument was used. 3C-like proteinase (Mybiosource), Biacore Amine Coupling Kit (Cytiva), Series S Sensor Chip CM5 (Cytiva), and PBS containing 9.2% DMSO at pH 7.4 was used as running buffer.

#### Immobilization pH Scouting

The best immobilization condition for Mpro on CM5 chip was determined by scouting of a 10 mM sodium acetate buffer at three different pH values, pH 4.0, 4.5 and 5.0. We determined that pH 4.0 was the optimal pH for immobilization.

#### Surface preparation with immobilization of main protease

Mpro was immobilized on CM5 Sensor Chip by activating the surface with 0.4 mol/L EDC / 0.1 mol/L NHS at a flow rate of 30 μL/min for 7 min. Mpro was dissolved in a 10 mM sodium acetate buffer, pH 4.0, to yield a 30 μg/mL solution. Following activation, Mpro solution was injected over the activated sensor chip surface at a flow rate of 30 μL/min for 6 min. Final ligand (Mpro) immobilization levels achieved were 11330 response units (RUs). The excess hydroxysuccinimidyl on the surface were deactivated with 1M ethanolamine hydrochloride, pH 8.5, for 7 min at a flow rate of 30 μL/min. The surface of a reference flow cell was activated with 0.4 mol/L EDC/0.1 mol/L NHS and then deactivated with 1 mol/L ethanolamine, with respective flow rates and times.

#### Analyte Injection

Before analyte injection, the sensor chip surface was conditioned with three 12.5 mM NaOH pulses of 30 sec followed by three start-up cycles, allowing the response to stabilize before analyte injection. We used 8 different percent of DMSO solutions; 10.0%, 9.8%, 9.6%, 9.4%, 9.2%, 9.0%, 8.8% and 8.6% in the solvent correction cycles to correct responses for variations in the bulk refractive index of the samples. Data were collected at a temperature of 25°C and 1 Hz rate. Montelukast was tested from lowest to highest concentration. During each sample cycle, analyte was injected for 60 sec at a flow rate of 30 μL/min. Dissociation period was monitored for 120 sec after analyte injection before regeneration with 12.5 mM NaOH for 30 sec at a flow rate of 30 μL/min to wash any remaining analyte from the sensor chip and wash flow with 50% DMSO before running the next sample.

#### 1:1 Steady-State Binding Affinity Analysis

Responses measured in the blank flow cell (control) were subtracted from the response measured in the active flow cell. The binding affinity (KD) of the interaction was obtained by plotting double-referenced binding responses versus Montelukast concentration and fitting the curve using a 1:1 steady-state affinity with constant Rmax interaction model.

### 3.4. Cell Viability Assay

#### Cell Culture Conditions

The human embryonic kidney (HEK293T CRL-11268) and *Cercopithecus aethiops* kidney (Vero E6, CRL1586), human lung adenocarcinoma (Calu-3, HTB-55; A549, CCL185) cell lines were purchased from American Type Culture Collection (ATCC, USA). They were cultured in Dulbecco’s Modified Eagle Medium (DMEM, p# 41965 Gibco) including 10% Fetal Bovine Serum (FBS, p# 10500 Gibco), 1% antibiotic-antimycotic in the tissue flasks and incubated at 37 ^o^C and 5% CO2. They were subcultured and used for cell viability and pseudovirus neutralization assay when they reach 70 - 80% confluency.

#### Cell Viability Assay

HEK293T, Vero E6, Calu-3, and A549 cells were cultured at a cell density of 1 x 10^4^ in the 96-well plates incubated at 37 ^o^C, and % 5 CO2 for 24 hrs. Following day, after aspiration of the medium, Montelukast (p# 1446859 Sigma-Aldrich) was added at concentrations of 1, 5, 10, 25, 50, 100, and 200 μM in DMEM and incubated for 24, 48, and 72 hrs. To measure the cell viability, MTT (3-[4,5-dimethylthiazol-2-yl]-2,5-diphenyltetrazolium bromide, p# M5655, Sigma-Aldrich) was dissolved in Phosphate Buffer Solution (PBS, 5 mg/ml). After each incubation periods, MTT at 5 μg/ml in DMEM was added into the each well and incubated at 37 °C and 5 % CO2 for 4 h. The formazan crystals dissolving solution, dimethyl sulfoxide (DMSO p# D 8418, Sigma-Aldrich) was added into the each well and incubated for 2 hrs. The absorbance was measured on an ELISA plate reader with a test wavelength of 570 nm and a reference wavelength of 630 nm.

#### 3.4.1 Pseudovirus production and infection

##### Transfection

HEK293T is highly transfectable cell line and widely used for retroviral production. Lentiviral-based pseudoviruses bearing SARS-CoV-2 Spike (S) or VSV-G glycoproteins were produced based on previous studies.^39^ Briefly, HEK293T cells were seeded at a cell density of 5×10^5^ cells/well on the 6-well plates. Next day, the cells in each well at approximately 70-80% confluency were used for transfection. After aspiration of the medium from each well, the transfection agent Fugene-6 in 10 μl was added on the 100 μl of DMEM basal medium (without FBS and Pen/Strep) in the 1.5 ml of tube and incubated 5 min at room temperature (RT). In another tube, 7500 ng of lenti RRL_GFP reporter plasmid, 6750 ng of psPAX2 packaging plasmid (Addgene plasmid # 12260), and 750 ng of Spike-18aa truncated (Addgene plasmid # 149541) were mixed. The plasmid mix was added into the fugen-6 tube and incubated 25-30 min at RT and then placed drop by drop over the cells in the well. After 14-16h of transfection, the media was removed, and fresh full media (DMEM with 10% FBS and 1% Pen/Strep) was added on the cells. After 48h of transfection, the pseudoviruses were collected, and filtered through 0.45 μm syringe filters, and stored at +4 °C for short term usage (up to 3-4 days), or stored at −80 ^o^C for long term storage.

#### 3.4.2 Pseudovirus Neutralization Assay

To render the cells infection by pseudoviruses, HEK293T cells on the wells were co-transfected with 1250 ng of ACE2 (Addgene plasmid #141185) and 1250 ng of TMPRSS2 expression plasmids (Addgene plasmid #145843) in the 6-well plate. After 48 hr of transfection period, HEK293T cells were harvested and seeded at 2×10^4^ cells/well on 96-well black plates and incubated at 37 °C and 5% CO2 for 24 hr. Following day, 50 μl previously prepared pseudoviruses and 50 μl of each tested concentration of Montelukast in the conditioned media were mixed in the tubes and incubated for 60 min at RT then directly used to infect ACE2 and TMPRSS2-expressing HEK293T cells. The infection rate by pseudoviruses was determined by measuring fluorescence intensity due to GFP reporter plasmids in the microplate reader. The cell viability at the same wells was determined by using CellTiter-Glo Luminescent Cell Viability Assay Kit (# G7571, Promega). Neutralization efficiency was calculated as relative fluorescence to the conditioned media collected from mock-transfected cells.

### 3.5. Virus Neutralization Assay Using xCELLigence MP Real Time Cell Analyzer

In the experiment, VERO E6 cell line (passage number: 17) was used. VERO E6 cells were cultured in Dulbecco’s Modified Eagle Medium with low glucose (DMEM/LOW GLUCOSE, HyClone, Cat # SH30021.01, lot # AF29484096) supplemented with final concentration of 10% heat-inactivated fetal bovine serum (FBS, HyClone, Cat # SV30160.03, lot # RE00000002) and 1% penicillin (10,000 Units/ml) - streptomycin (10,000 Units/ml) (HyClone, Cat # SV30010, Lot # J190007). Sample stocks were diluted in DMEM low glucose supplemented with 2% FBS to make a concentration range. Neutralization assay was performed based on impedance using xCELLigence MP real time cell analyzer equipment (RTCA). The impedance is expressed as arbitrary units called cell index. VERO E6 cells were suspended in DMEM low glucose supplemented with 10% FBS and seeded into a disposable sterile 96-well E-plate of the xCELLigence RTCA MP device as final cell concentration of 2.5×10^4^ cells per well. The instrument has been placed in a CO_2_ cell culture incubator during the experiment, and controlled by a cable connected to the control unit located outside the incubator. The cells on 96-well E-Plate was placed in the xCELLigence RTCA MP device, and incubated for 24 hours. After incubation, the cells were inoculated with 10^6^ PFU SARS-CoV2 virus for an hour in DMEM low glucose supplemented with 2% FBS. Then, the Montelukast samples that were prepared in accordance with the sample preparation protocol in different concentrations (375 μM, 188 μM, 94 μM, 47 μM, 23 μM, 12 μM, 6 μM, 3 μM and 1.5 μM) were added to the wells of the E-Plate. Cells were incubated in 37°C, 5% CO2 humidified incubator. Real time measurement for each cell line was collected for 150 hours with intervals of 15 min. At the end of the period, the experiment was terminated and the data obtained were analyzed using RTCA Software Pro software.

The electrical conductivity is converted into the unitless cell index (CI) parameter by the xCELLigence MP device every 15 minutes. The increase in CI indicates cell viability / health, whereas a decrease indicates cell death / unhealth. It is expected that VERO E6 cells in the presence of the SARS-CoV2 virus will demonstrate a decrease in CI values. The data obtained in the graphic in figures are normalized according to the time point when the virus was added to the experiment. The samples in the figure are indicated according to their color codes.

## 4. Conclusions

Here, the potential effect of the Montelukast on SARS-CoV-2 is investigated using multiscale molecular modeling approaches and integrated *in vitro* experiments including FRET-based binding assays, SPR, pseudovirus and virus neutralization methods. Our results show that Montelukast has dual inhibitor effect and exerts its effect on SARS-CoV-2 by interference with the entry of the virus into the host cell (via Spike/ACE-2) as well as it inhibits the 3C-like protease which is responsible for functional protein maturation. Our results show that 12 μM Montelukast concentration delays the CIT_50_ time by about 12 hours. It has been observed that Favipiravir, which is one of the mostly used antiviral drugs in COVID-19 therapy, should be used in 16-fold more doses than Montelukast in order to show the same effect. Since Montelukast is an approved drug and has been widely used in the market for over 20 years against asthma, its side effects have been well studied and the results show that it is a well-tolerated drug. Since its patent is expired in 2012, its clinical usage at COVID19 can be urgently considered.

## Acknowledgment

This study was funded by Scientific Research Projects Commission of Bahçeşehir University. Project number: BAU.BAP.2020.01. This study was also funded by the The Scientific and Technological Research Council of Turkey (TÜBİTAK), within the program number of 18AG003. We would like to thank Néstor Santiago-Gonzalvo (Cytiva) for his helpful discussion and support with the Biacore experiments.

